# Single Cell RNA Sequencing of stem cell-derived retinal ganglion cells

**DOI:** 10.1101/191395

**Authors:** Maciej Daniszewski, Anne Senabouth, Quan Nguyen, Duncan E. Crombie, Samuel W. Lukowski, Tejal Kulkarni, Donald J Zack, Alice Pébay, Joseph E. Powell, Alex W. Hewitt

## Abstract

We used human embryonic stem cell-derived retinal ganglion cells (RGCs) to characterize the transcriptome of 1,174 cells at the single cell level. The human embryonic stem cell line BRN3B-mCherry A81-H7 was differentiated to RGCs using a guided differentiation approach. Cells were harvested at day 36 and subsequently prepared for single cell RNA sequencing. Our data indicates the presence of three distinct subpopulations of cells, with various degrees of maturity. One cluster of 288 cells upregulated genes involved in axon guidance together with semaphorin interactions, cell-extracellular matrix interactions and ECM proteoglycans, suggestive of a more mature phenotype.

## BACKGROUND & SUMMARY

Since the isolation of the embryonic stem cells (ESCs)^1–4^ and generation of induced pluripotent stem cells (iPSCs)^5,6^, pluripotent stem cells (PSCs) have made a tremendous contribution towards improving our understanding of mechanisms involved in development and disease. PSCs have the ability to self-renew and differentiate into all cell types of the body, thereby providing great potential for regenerative medicine and cell replacement therapies. Further, PSC-derived progeny allow investigating disease-affected cell types that are not readily accessible due to their anatomical location, such as retinal cells ^7–9^. Utilising such disease-affected cells will also significantly improve the drug development pipeline through efficacy profiling and side effect or toxicity assessment ^10^.

The development of RNA sequencing (RNA-seq) technology has allowed for the rapid quantification of individual gene transcripts. Integrating this high-throughout data with computational and statistical methods provides a toolbox to study the molecular functions of human tissues. Critically, to date, the majority of RNA-seq studies have been conducted on ‘bulk’ samples, consisting of millions of individual cells - the result of which is that transcript quantification represents the average signal across that cell population. Recent developments to isolate single cells, and barcode their expressed transcripts has enabled the transcriptomes of single cells to be sequenced (scRNA-seq) in a high-throughput manner. By sequencing large number of single cells from an individual ‘sample’ it is now possible to dissect the cellular composition of apparently homogenous tissues or cell culture ^11–13^. scRNAseq also opens the possibility of examining rare cell populations that could not otherwise be resolved using bulk RNA-seq, and further characterising well-known cell types, for example oligodendrocytes ^14^ or sensory neurons ^15^. Moreover, scRNA-seq may also be used for tracking cell development during differentiation, as movement between different cell types is associated with changes in gene expression. Thus, stages across a cell lineage can be distinguished by their unique transcriptional signature ^16^. This technology has also been used in cell culture, in particular with PSCs, their differentiated progeny and organoids, including of the nervous system ^17,18^, as a way to distinctively characterize cellular subpopulations. Results of such analyses can discern determinants of cell fates, and this information can then applied to *in vitro* differentiation experiments to increase efficiency of generating the tissue of interest ^19^.

Retinal ganglion cells (RGCs) transmit pre-processed signals from the retina to the midbrain through the optic nerve. Many diseases, such as primary open angle glaucoma, Leber hereditary optic neuropathy and autosomal dominant optic atrophy manifest by degeneration or loss of RGCs and culminate in irreversible loss of sight. It is estimated that there are more than 30 subtypes of RGCs in mammalian retina ^20,21^; however, the molecular profiling of RGCs in human disease has proven difficult. Currently, studying optic neuropathies is hindered by the lack of non-invasive means for obtaining RGCs from living donors. This can now be circumvented by use of PSCs as a source of RGCs ^8^. We recently described a protocol for the differentiation of human PSCs into functional RGCs ^7^. RGCs generated through this method are functional, as exemplified by the presence of sodium and potassium currents, mature axon potentials and the expression of RGC-specific markers, including *BRN3B*, *ISL1* and *PRPH* ^7^. Moreover, whole transcriptome analysis through bulk RNA-seq of our PSC-derived RGCs demonstrated close resemblance to sensory neurons, and cells from the ganglion cell layer ^7^. Herein we present a dataset of scRNA-seq to enable the characterization of the transcriptome of RGCs derived from human ESCs (hESCs) at a single cell level.

## METHODS

### Ethical Approval

All experimental work performed in this study was approved by the Human Research Ethics committees of the University of Melbourne (0605017) with the requirements of the National Health & Medical Research Council of Australia (NHMRC) and conformed with the Declarations of Helsinki.

### Cell culture and retinal differentiation

The reporter line BRN3B-mCherry A81-H7 hESC^9^ was maintained on vitronectin-coated 6-well plates using StemFlex (Gibco). Culture medium was changed every second day. Cells were differentiated into RGCs as previously described ^7^. Briefly, undifferentiated hESCs cultured in monolayer on vitronectin-coated plates were differentiated using RGC differentiation medium 2 (DMEM F12 with GlutaMAX, 10% KnockOut Serum Replacement (Invitrogen), SM1 (Stem Cell Tech), 10 ng/mL noggin (Sapphire Biosciences), 10 ng/ml Dickkopf-related protein 1 (DKK1, Peprotech), 10 ng/ml Insulin Growth Factor 1 (IGF1, Peprotech) and 5 ng/ml basic Fibroblast Growth Factor (bFGF, Merck). Medium was changed every 2-3 days. RGC differentiation was monitored by the appearance of mCherry-positive cells, reflective of *BRN3B* expression.

### Fluorescence-activated cell sorting (FACS)

On day 36 of differentiation, cells were washed with phosphate-buffered saline (PBS) and incubated with Accutase (Sigma, 37°C, 5 minutes). Cells were then incubated in RGC differentiation medium supplemented with the ROCK inhibitor Y27632 (10 μM, Selleckchem, RGC+RI) and gently dissociated using a P1000 pipette, filtered using a 100 μm nylon strainer (BD Falcon) and centrifuged (300g, 10 minutes). The cell pellet was resuspended in RGC+RI medium and incubated with THY1 antibody (Miltenyi, 4°C, 15 minutes). Cells were washed in RGC+RI medium, and centrifuged (300g, 3 minutes). Two modifications to our original protocol were performed. Firstly, selection of RGCs using THY1 was performed by FACS instead of the magnetic sorting we originally reported. Secondly, cells were prepared for sequencing immediately following THY1 selection and were not allowed to rest prior to being further processed. A cell pellet was resuspended in 500 μl of RGC+RI prior to sorting with a BD FACSAria III cell sorter (Becton, Dickinson). Both THY1-positive (+ve) and THY1-negative (-ve) fractions were collected in 5 mL conical tubes (BD Falcon).

### Single-cell preparation

Both THY1-positive (+ve) and THY1-negative (-ve) fractions were subjected to library preparation using the Single Cell 3’ Reagent Kit (10X Genomics) as per the manufacturer’s instruction. This step was performed within 60 minutes of the FACS. Briefly, cell suspension was mixed using a wide-bore tip to determine cell concentration using a Countess® Automated Cell Counter (Life Technologies). Cells were centrifuged for 5 minutes at 300g and cell pellet was resuspended in PBS with 0.04% BSA. Cell suspension was passed through a cell strainer to remove any remaining cell debris and large clumps and cell concentration was determined again.

### Generation of single cell GEMs and sequencing libraries

Single cell suspensions were loaded onto 10X Genomics Single Cell 3' Chips along with the reverse transcription (RT) master mix as per the manufacturer's protocol for the Chromium Single Cell 3' v2 Library (10X Genomics; PN-120233), to generate single cell gel beads in emulsion (GEMs). Sequencing libraries were generated with unique sample indices (SI) for each sample. The resulting libraries were assessed by gel electrophoresis (Agilent D1000 ScreenTape Assay) and quantified with qPCR (Illumina KAPA Library Quantification Kit). Following normalization to 2nM, libraries were denatured and diluted to 17pM of cluster generation using the Illumina cBot (HiSeq PE Cluster Kit v4). Libraries for the two samples were multiplexed respectively, and sequenced on an Illumina HiSeq 2500 (control software v2.2.68/ Real Time Analysis v1.18.66.3) using a HighSeq SBS Kit v4 (Illumina, FC-401-4003) in high-output mode as follows: 126bp (Read 1), 8bp (i7 Index), 8bp (i5 Index), and 126bp (Read 2).

### Mapping of reads to transcripts and cells

The sequencing data was processed into transcript count tables with the Cell Ranger Single Cell Software Suite 1.3.1 by 10X Genomics (http://10xgenomics.com/). Raw base call files from the HiSeq4000 sequencer were demultiplexed with the cellranger mkfastq pipeline into library-specific FASTQ files. As the libraries were sequenced using non-standard settings, cellranger mkfastq was run with the following parameters: --use-bases-mask="Y26n*,I8n*,n*,Y98n*" --ignore-dual-index. The FASTQ files for each library were then processed independently with the cellranger count pipeline. This pipeline used STAR ^22^ to align cDNA reads to the Homo sapiens transcriptome (Sequence: GRCh38, Annotation: Gencode v25). Once aligned, barcodes associated with these reads – cell identifiers and Unique Molecular Identifiers (UMIs), underwent filtering and correction. Reads associated with retained barcodes were quantified and used to build a transcript count table. Resulting data for each sample were then aggregated using the cellranger aggr pipeline, which performed a between-sample normalization step and concatenated the two transcript count tables. Post-aggregation, the mapped data was processed and analyzed as described below.

### Preprocessing

To preprocess the mapped data, we constructed a cell quality matrix based on the following data types: library size (total mapped reads), total number of genes detected, percent of reads mapped to mitochondrial genes, and percent of reads mapped to ribosomal genes. Cells that had any of the 4 parameter measurements higher than 3x median absolute deviation (MAD) of all cells were considered outliers and removed from subsequent analysis (Table 1). In addition, we applied two thresholds to remove cells with mitochondrial reads above 20% or ribosomal reads above 50% (Table 1). To exclude genes that were potentially detected from random noise, we removed genes that were detected in fewer than 1% of all cells.

**Table 1.** Summary statistics for sequencing and mapping data of two samples

| Sample | Number of cells | Median reads per cell | Median genes per cell | Total genes detected | Median UMIs per cell | Total number of reads | Percent mapped reads | Remaining cells post filtering |
| --- | --- | --- | --- | --- | --- | --- | --- | --- |
| 1 | 1,090 | 124,127 | 3,528 | 21,317 | 13,575 | 135,299,096 | 61.10 | 993 |
| 2 | 194 | 493,659 | 5,188 | 19,812 | 26,218 | 95,769,921 | 62.80 | 181 |

The expression data was normalised on two levels to reduce possible systematic bias between samples and between cells. The first level of normalisation - between samples, was performed prior to data aggregation using cellranger aggr’s depth equalisation method ^23^. This method reduces potential confounding effects caused by differences in sequencing depths between samples by subsampling mapped reads from higher-depth libraries until the number of mapped reads per library were equal. The second level of normalisation - between cells, was performed after filtering using the deconvolution approach by Lun *et al*. ^24^. This level of normalisation reduces bias possibly caused by technical variation such as cDNA synthesis, PCR amplification efficiency and sequencing depth for each cell. The deconvolution approach was chosen as it accounts for the sparse nature of expression data by pooling expression counts from groups of cells. As the sizes of the groups were linear (40, 60, 80, 100), the group-specific normalised size factors could be deconvolved into cell-specific size factors that were then used to scale the counts of individual cells. After normalisation, abundantly expressed ribosomal protein genes and mitochondrial genes were discarded. We have made available both the raw and normalised data on ArrayExpress under accession E-MTAB-6108.

### Identification of residual low-quality cells via clustering

We identified and removed a small group of cells with low-quality sequence data. These cells were not detected by initial filtering; instead, they were identified via clustering and enrichment of differentially expressed genes. The transcript count table underwent dimensionality reduction using Principal Component Analysis (PCA). This procedure was applied to the top 1,500 most variable genes using the *prcomp()* function in R ^25^. The first 20 PCs were retained and a cell-PCA eigenvector matrix was used for clustering.

We applied an unsupervised clustering method that does not take into account any predetermined parameters to objectively identify single cell subpopulations ^26^. This method is less biased compared to top-down clustering approaches, such as the k-means. Briefly, to achieve high-resolution clustering capable of detecting small subpopulations and outliers, we applied bottom-up agglomerative hierarchical clustering to construct a dendrogram tree, which contains one cell as one bottom branch (the highest resolution). We used the reduced dataset containing the top 20 PCs described above to calculate an Euclidean distance matrix between cells, and organized cells into the dendrogram using the Ward’s minimum distance so that similar cells are joint into larger groups of branches. To identify subpopulations, we applied an unsupervised, objective approach to merge the branches into a high-resolution and stable clustering result. The approach divided the dendrogram tree into 40 height-windows, ranging from 0.025 (from the bottom of the tree) to 1 (from the top). By iteratively and dynamically merging cells in each of the 40 height-windows, we generated 40 independent clustering results, different in the clustering resolutions. Clustering results were then compared quantitatively using adjusted rand indices, which score pairs of cells that are the same or different between two clustering results ^27^. The optimal clustering result was the most stable result across a range of consecutive tree-height values.

To characterise the identified clusters, we performed pairwise differential expression analysis by fitting a general linear model and using a negative binomial test as described in the DESeq package ^28^. Network analysis was then performed on significant differentially-expressed genes using Reactome functional interaction analysis ^28,29^.

### Code availability

All code and usage notes available at: https://github.com/IMB-Computational-Genomics-Lab/RetinaGanglionCells. This includes: computational bioinformatic pipelines that process sequence data in BCL format through to a mapped UMI expression matrix; Scripts for quality-control, normalisation, clustering, differential expression and visualization.

### Data Records

Data is available at ArrayExpress under accession number: E-MTAB-6108. Files consist of raw FASTQ files as well as a tab separated matrix of Transcripts Per Million for each cell passing quality control filtering.

## TECHNICAL VALIDATION

The hESC reporter line BRN3B-mCherry A81-H7 was differentiated to RGCs following our established protocol ^7^, changing culture medium every second day. After 36 days, selection of RGCs using THY1 was performed by FACS. Both positive and negative THY fractions were harvested, and single cells harvested for library preparation and scRNA-Seq as outlined in Figure 1. Processing our initial analysis identified a group of 61 cells whose expression levels indicated degradation and apoptosis (Figure 2). These 61 cells were removed from the data and the expression data from the remaining 1,174 healthy cells was re-normalised and analysed. Clustering of these 1,174 cells identified three distinct subpopulations consisting of 531, 355 and 288 cells (Figure 3A). We performed differential expression analysis and subsequently pathway enrichment to characterise the molecular functions of these subpopulations (Figure 3B).

**Figure 1:**
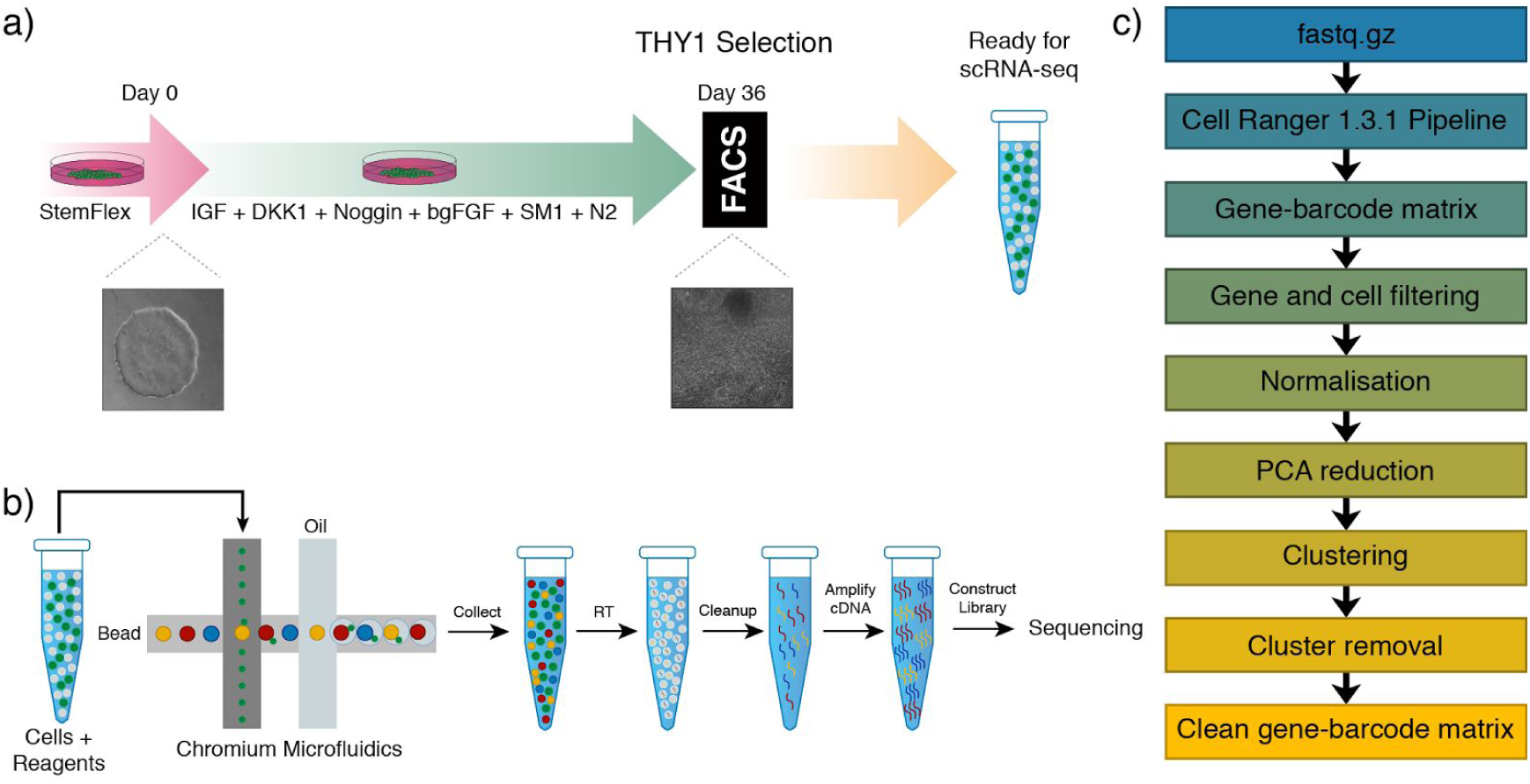
Schematic representation of the experimental workflow. (**a**) Guided differentiation of the reporter line BRN3B-mCherry A81-H7 hESCs into RGCs using IGF1, DKK1, Noggin, bFGF in a neural medium containing SM1 and N2, as described in [28]. On day (d) 36, cells were sorted based on the expression of the marker THY1. Cells from both positive and negative THY fractions were then processed for scRNA-seq. Brightfield images describe cell morphology of undifferentiated hESCs prior to differentiation (d0) and post differentiation (d36) at time of sorting. (**b**) Single cell suspensions are prepared and libraries generated using the Chromium V2 chemistry. Libraries were sequenced on an Illumina HiSeq2500. **(c)** Sequence data is processed using bioinformatic pipelines, and analysis conducted on the resulting expression matrix.

**Figure 2:**
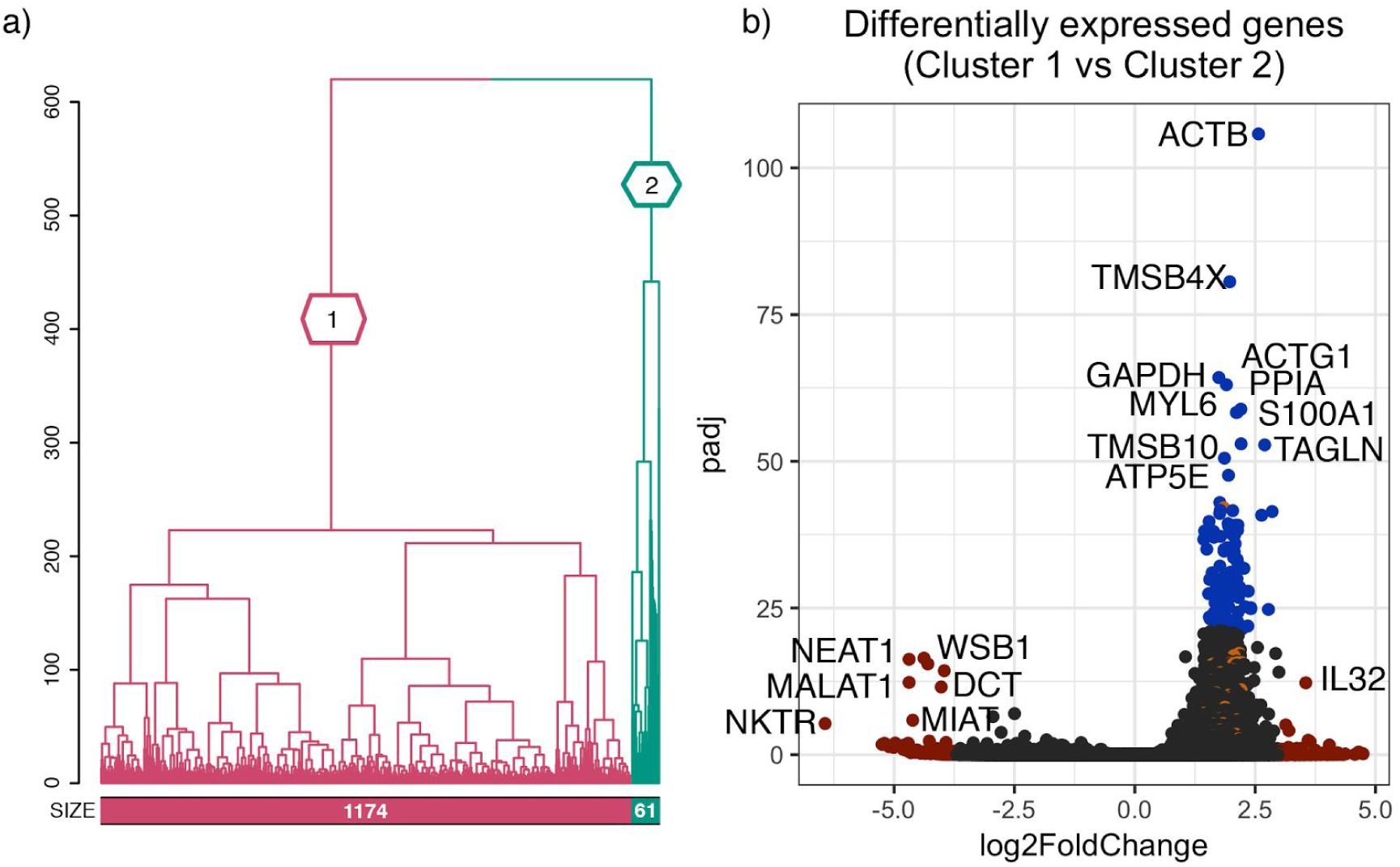
Identification of residual low-quality cells via clustering. **(a)** Unsupervised clustering of all cells into two subpopulations. The dendrogram tree displays distance and agglomerative clustering of the cells. Each branch represents one subpopulation. The clustering is based on the most stable clustering result across 40 tree cut heights. The branches are labelled with their subpopulation identification. The number of cells in each of the two populations are given below the branches. **(b)** The top significantly expressed genes of cells in subpopulation one vs two. Genes represented by blue and red points are those in the top 0.5% highest log2 fold-change or -log10(*p*-value) respectively. Genes represented by orange points are related to apoptotic pathways.

**Figure 3:**
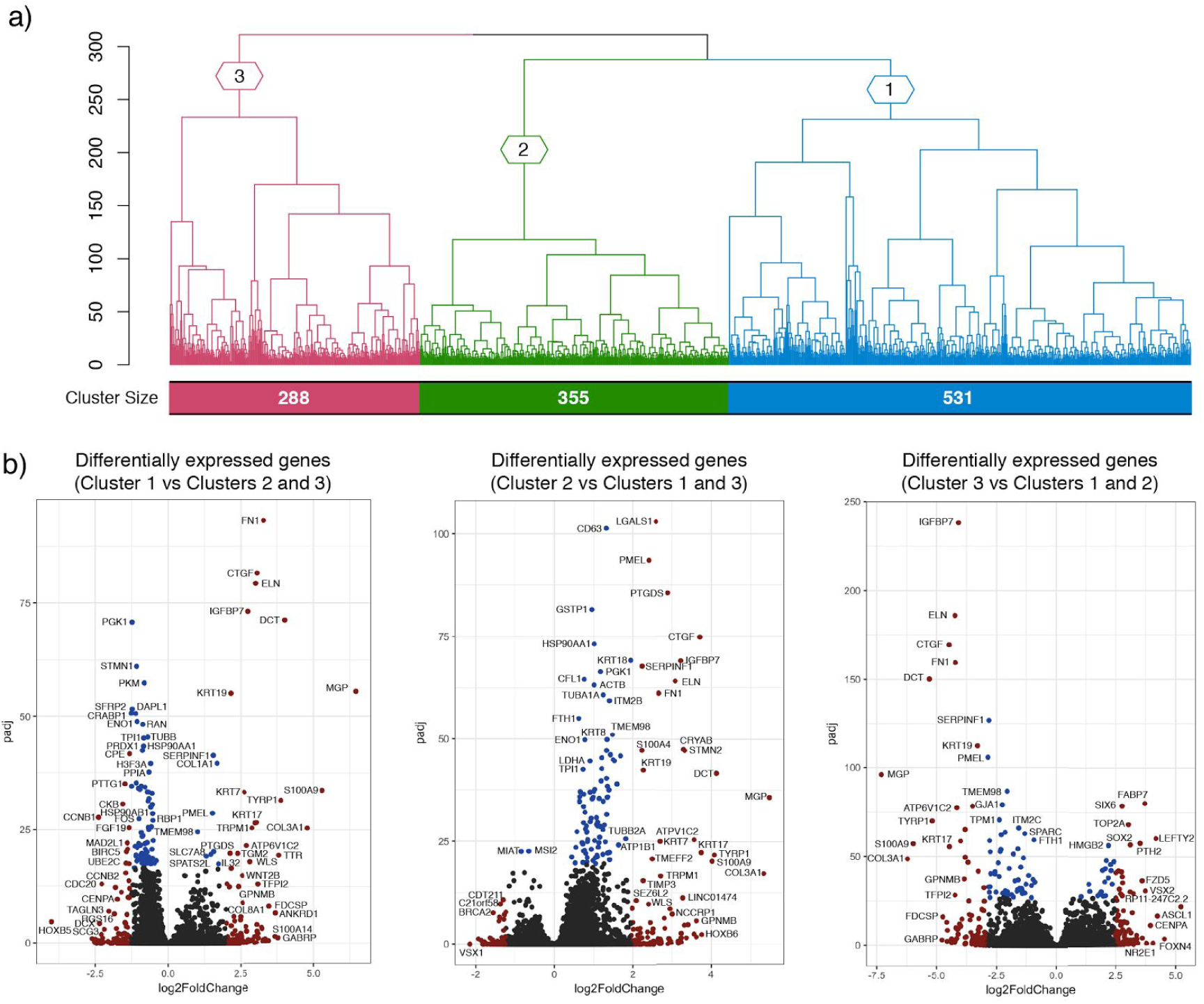
Characterisation of filtered cells via clustering and differential expression. **(a)** Unsupervised clustering of cells after filtering into three subpopulations. The dendrogram tree displays distance and agglomerative clustering of the cells. Each branch represents one subpopulation. The clustering is based on the most stable clustering result across 40 tree cut heights. The branches are labelled with their subpopulation identification. The number of cells in each of the three populations are given below the branches. **(b)** The top significantly expressed genes of cells of each cluster vs other clusters. Genes represented in blue and red points are those in the top 0.5% highest logFC or -log(p-value) respectively.

The 531 cells from subpopulation one were upregulated for genes associated with neural cell adhesion molecule signaling for neuronal outgrowth and Hedgehog pathway, which plays various roles in patterning of the central nervous system. Interestingly, genes implicated in collagen biosynthesis, extracellular matrix proteoglycans and axon guidance were downregulated (**Table S1**). This pattern of gene expression suggests a progenitor or an early differentiation state. Cells from subpopulation two contained upregulated genes associated with control of the Notch protein expression, which is a crucial member of the Notch signaling pathway implicated in the neuronal function and development, and DNA repair (**Table S2**). Collectively, this pattern of gene expression is indicative of a more differentiated RGC phenotype than the cells in cluster one. The 288 cells identified as subpopulation three contained upregulated genes involved in axon guidance, together with semaphorin interactions, cell-extracellular matrix interactions and extracellular matrix proteoglycans. Furthermore, we observed significant downregulation of multiple genes associated with cell cycle (**Table S3**). Taken together this indicates that this subpopulation three represents a more mature neuronal phenotype compared to cells in the other two subpopulations. Importantly, one cell within subpopulation one was found to express *OPN4* a gene known to be expressed in intrinsically-photosensitive RGCs. Thus, these data indicate different levels of maturity of ESC derived RGC, with this conclusion supported by observed pathway enrichment (**Table S4-6**).

### Usage Notes

Our experiment was designed to assess the different subpopulations of RGCs post differentiation from hESCs. hESC-derived RGCs obtained in a 36 day guided differentiation clustered into distinct subpopulations of neurons. Our initial analysis identified a group of 61 cells that showed a strong enrichment of stress and apoptosis pathways. This is possibly due to the FACS procedure itself, which can be stressful on cells. All post quality-control cells express genes relevant to RGC structure and functions. Altogether, our data provides strong support of an RGC identity of the cells in all clusters.

## ACKNOWLEDGEMENTS

This work was supported by grants from the Sir Edward Dunlop Medical Research Foundation, the Ophthalmic Research Institute of Australia, Retina Australia, the Joan and Peter Clemenger Foundation, and a Philip Neal bequest. AWH & JEP are supported by National Health and Medical Research Council Fellowships, AP by an Australian Research Council Future Fellowship (FT140100047), and MD by an International Postgraduate Award Scholarship from the University of Melbourne. CERA receives Operational Infrastructure Support from the Victorian Government.

## AUTHOR CONTRIBUTIONS

MD: concept and design, experimental work, interpretation of data, writing of manuscript, final approval of manuscript.

AS: concept and design, experimental work, interpretation of data, writing of manuscript, final approval of manuscript.

QN: experimental work, interpretation of data, writing of manuscript, final approval of manuscript.

DEC: interpretation of data, final approval of manuscript. SWL: experimental work, final approval of manuscript. TJ: interpretation of data, final approval of manuscript.

DJZ: provision of reagents, final approval of manuscript.

AP: concept and design, financial support, interpretation of data, writing of manuscript, final approval of manuscript.

JEP: concept and design, financial support, interpretation of data, writing of manuscript, final approval of manuscript.

AWH: concept and design, financial support, interpretation of data, writing of manuscript, final approval of manuscript.

## REFERENCES

1. Martin, G. R. Isolation of a pluripotent cell line from early mouse embryos cultured in medium conditioned by teratocarcinoma stem cells. Proc. Natl. Acad. Sci. U. S. A. 78,7634–7638 (1981).

2. Evans, M. J. & Kaufman, M. H. Establishment in culture of pluripotential cells from mouse embryos. Nature 292, 154–156 (1981).

3. Reubinoff, B. E., Pera, M. F., Fong, C. Y., Trounson, A. & Bongso, A. Embryonic stem cell lines from human blastocysts: somatic differentiation in vitro. Nat. Biotechnol. 18,399–404 (2000).

4. Thomson, J. A. Embryonic Stem Cell Lines Derived from Human Blastocysts. Science 282, 1145–1147 (1998).

5. Takahashi, K. & Yamanaka, S. Induction of pluripotent stem cells from mouse embryonic and adult fibroblast cultures by defined factors. Cell 126, 663–676 (2006).

6. Takahashi, K. et al. Induction of pluripotent stem cells from adult human fibroblasts by defined factors. Cell 131, 861–872 (2007).

7. Gill, K. P. et al. Enriched retinal ganglion cells derived from human embryonic stem cells. Sci. Rep. 6, 30552 (2016).

8. Gill, K. P., Hewitt, A. W., Davidson, K. C., Pébay, A. & Wong, R. C. B. Methods of Retinal Ganglion Cell Differentiation From Pluripotent Stem Cells. Transl. Vis. Sci. Technol. 3, 7 (2014).

9. Sluch, V. M. et al. Differentiation of human ESCs to retinal ganglion cells using a CRISPR engineered reporter cell line. Sci. Rep. 5, 16595 (2015).

10. Cook, D. et al. Lessons learned from the fate of AstraZeneca’s drug pipeline: a five-dimensional framework. Nat. Rev. Drug Discov. 13, 419–431 (2014).

11. Shalek, A. K. et al. Single-cell transcriptomics reveals bimodality in expression and splicing in immune cells. Nature 498, 236–240 (2013).

12. Wills, Q. F. et al. Single-cell gene expression analysis reveals genetic associations masked in whole-tissue experiments. Nat. Biotechnol. 31, 748–752 (2013).

13. Kolodziejczyk, A. A. et al. Single Cell RNA-Sequencing of Pluripotent States Unlocks Modular Transcriptional Variation. Cell Stem Cell 17, 471–485 (2015).

14. Marques, S. et al. Oligodendrocyte heterogeneity in the mouse juvenile and adult central nervous system. Science 352, 1326–1329 (2016).

15. Usoskin, D. et al. Unbiased classification of sensory neuron types by large-scale single-cell RNA sequencing. Nat. Neurosci. 18, 145–153 (2015).

16. Etzrodt, M., Endele, M. & Schroeder, T. Quantitative single-cell approaches to stem cell research. Cell Stem Cell 15, 546–558 (2014).

17. Quadrato, G. et al. Cell diversity and network dynamics in photosensitive human brain organoids. Nature 545, 48–53 (2017).

18. Birey, F. et al. Assembly of functionally integrated human forebrain spheroids. Nature 545, 54–59 (2017).

19. Tang, F. et al. Tracing the derivation of embryonic stem cells from the inner cell mass by single-cell RNA-Seq analysis. Cell Stem Cell 6, 468–478 (2010).

20. Rousso, D. L. et al. Two Pairs of ON and OFF Retinal Ganglion Cells Are Defined by Intersectional Patterns of Transcription Factor Expression. Cell Rep. 15, 1930–1944(2016).

21. Macaulay, I. C. et al. G&T-seq: parallel sequencing of single-cell genomes and transcriptomes. Nat. Methods 12, 519–522 (2015).

22. Dobin, A. et al. STAR: ultrafast universal RNA-seq aligner. Bioinformatics 29, 15–21(2013).

23. Zheng, G. X. Y. et al. Massively parallel digital transcriptional profiling of single cells. Nat. Commun. 8, 14049 (2017).

24. Lun, A. T. L., Bach, K. & Marioni, J. C. Pooling across cells to normalize single-cell RNA sequencing data with many zero counts. Genome Biol. 17, 75 (2016).

25. McCarthy, D. J., Campbell, K. R., Lun, A. T. L. & Wills, Q. F. Scater: pre-processing, quality control, normalization and visualization of single-cell RNA-seq data in R. Bioinformatics 33, 1179–1186 (2017).

26. Nguyen, Q. et al. Single-Cell Transcriptome Sequencing Of 18,787 Human Induced Pluripotent Stem Cells Identifies Differentially Primed Subpopulations. (2017).doi:10.1101/119255

27. Rand, W. R. Objective Criteria for the Evaluation of Clustering Methods. J. Am. Stat. Assoc. 66, 846–850 (1971).

28. Anders, S. & Huber, W. Differential expression analysis for sequence count data. Genome Biol. 11, R106 (2010).

29. Wu, G., Feng, X. & Stein, L. A human functional protein interaction network and its application to cancer data analysis. Genome Biol. 11, R53 (2010).

